# Production of green synthesized iron oxide /graphene magnetic nanocomposite as a novel antibacterial material

**DOI:** 10.1101/2024.09.12.612624

**Authors:** Mohammad kazazi, Ali Moomivand, Ali Mahmoodvand

## Abstract

Synthesis of green iron magnetic nanoparticles (Nps) based on graphene is an environmentally friendly method in nanotechnology. In this method, graphene is synthesized via electrochemical process from graphite. Then, a homogeneous solution of graphene in a mixture of water and alcohol is prepared, and iron magnetic Nps are synthesized using iron(II) chloride and iron(III) chloride in a 1:2 ratio. Finally, the antibacterial property of the synthesized nanocomposite was investigated, and analyses including SEM, XRD, FT-IR, EDAX, and VSM were performed.

---

In the past three decades, due to the increase in problems caused by chemical pollutants and the reduction of available resources, green chemistry methods have emerged. The objectives of green chemistry include improving efficiency, reducing process steps, using safer chemicals, and use less hazardous raw materials [1]. Moreover, in recent years, due to the significant benefits of green chemistry, efficient green chemical methods for synthesizing metal Nps have gained the attention of many researchers. There have been considerable efforts to promote the use of green chemistry methods and environmentally compatible materials in Nps synthesis, such as utilizing biological organisms like microorganisms, plant extracts, or biomass as alternatives to chemical and physical methods for environmentally friendly Nps production [2-5].

Among the various common methods for producing Nps, the use of plant extracts has shown positive results. Generally, plant extracts, which act as stabilizers and coating agents for controlling crystal growth, consist of various metabolites such as terpenoids, phenols, proteins, or carbohydrates. These compounds directly enable the extract’s ability to perform redox reactions on Nps. Environmental pollutants, such as chemical and physical contaminants, can arise from the various chemical and physical processes used in Nps production. Therefore, researchers have made considerable efforts to synthesize nanomaterials using environmentally friendly processes with plant extracts, aiming for simpler and safer research frameworks [6, 7].

Nowadays, magnetic Nps have found diverse applications due to features like large specific surface area and easy separation using an external magnetic field. In food materials, magnetic Nps can be used for enzyme immobilization, protein purification, and analyzing relevant compositions.

In summary, magnetic Nps contain magnetic elements like iron, cobalt, nickel, and their chemical compounds. Regarding the application of magnetic Nps in food materials, assessing the safety or toxicity of these particles is of utmost importance. Among various types of magnetic Nps, iron oxide Nps, especially superparamagnetic Fe_3_O_4_ Nps, have found the most applications in the food sector due to their non-toxicity and good biological compatibility [8-12].

Traditional antibiotics based on organic components are traditionally used to treat infectious diseases originating from social and hospital environments. However, due to prolonged exposure of bacteria to antibiotics, they have developed resistance through various mechanisms such as gene mutation, against most commonly used antibiotics. Therefore, due to the overuse of antibiotics and increasing bacterial resistance, finding suitable alternatives to antibiotics is essential. For this reason, extensive studies have been conducted on the potential use of antimicrobial compounds found in plants, as well as the use of Nps to control and treat pathogens. Metal oxide Nps, due to their effective antibacterial activity, chemical stability, and relatively non-toxic properties, may act as efficient disinfectants [12-16]. Nanotechnology refers to the design, characterization, production, and use of structures and tools with control over shape and size at the nanometer scale (1-100 nm) [17]. The use of metals as antimicrobial and antibacterial agents has been prevalent since ancient times. However, with the emergence of modern and combination antibiotics, which have shown greater and faster effectiveness, and considering the destructive side effects of metals and their compounds on living tissues, they had been largely forgotten for years [18, 19].

In recent years, astonishing advances in nanoscience and the subsequent discovery of new properties of materials at the nanoscale have led researchers to explore metallic Nps as key solutions to the growing resistance problem of pathogenic microorganisms. Besides their amazing properties and high potential in biomedical applications, metallic Nps do not pose the harmful effects of metals and their ions in bulk dimensions on human health. Superparamagnetic iron oxide Nps exhibit distinct magnetic behavior, making them suitable for various aspects of cancer treatment, including imaging, thermal therapy, and targeted drug delivery [20-24].

The completely different mechanism of antibacterial activity of Nps compared to conventional antibiotics can be considered as a desirable and highly potential alternative for investigation and use. Recently introduced into scientific and industrial domains, new Nps such as graphene have emerged [25-27]. Reports indicate that due to the biophysical interaction between Nps and bacteria, they cause damage to the bacterial cell wall and penetrate the cell, leading to bacterial death [28-33].

During the investigated research, aqueous extract of green tea plant has been used to prepare supermagnetic Nps. This plant, with its specific medicinal properties attributed to iron and graphene, imparts antibacterial characteristics. Green tea, scientifically known as Camellia Sinensis, is a shrub or small tree reaching up to 9 m in height, featuring green leaves without lobes, obtained from the full-grown bud of Camellia Sinensis [34-36].

The proposed method in this study eliminates the use of chemical substances and processes for synthesizing iron and graphene magnetic Nps. This nanocomposite was utilized for dye removal and its antibacterial properties, and analyses including SEM, XRD, FT-IR, EDAX, and VSM were performed.

## Materials and Methods

### Materials

Iron (II) Chloride Hexahydrate, Iron (III) Chloride Tetrahydrate were purchased from the Merck Company, Green Tea was purchased from Touyserkan, Iran, Graphite Coal and Distilled Water.

### Synthesis of Graphene Nanoparticles

First, two medium-sized flashlight batteries are selected, then extracted graphite and placed in an electrolyte solution. They were connected to a direct current power source, one of the graphite rods acting as a working electrode and the other being consumed. The solution was evaporated and the remaining material was dried by an oven.

### Synthesis of Iron Oxide /Graphene Magnetic Nanocomposite

Initially, 1 g of graphene Nps was added to 100 ml of distilled water. The mixture was stirred for 30 min at 40 °C to ensure homogeneity. Subsequently, 1 g of iron (II) chloride hexahydrate and 2 g of iron (III) chloride tetrahydrate were added to the mixture and stirred for 30 min. Then, 100 ml of clarified green tea extract was added drop wise to the stirring solution, causing the solution to change color.

### Cultivation of Antibacterial of Iron Oxide /Graphene Magnetic Nanocomposite

The bacterium used for the antibacterial experiment was Escherichia coli (ATCC 25922) (E.coli). This bacterium was obtained from the Laboratory of Shahid Sardar Soleimani Hospital in Touyserkan County. A bacterial sample was cultured on specific Bald Agar medium and incubated for 24 h at 37 °C.

### Characteristics

We evaluated the microstructure, morphology, and chemical compound of the nanocomposite with a Field Emission Scanning Electron Microscope with Energy Dispersive X-Ray Spectroscopy(FE-SEM-EDAX) (Zeiss Sigma 300). The X-Ray Diffraction (XRD) patterns at angles 2θ= 32–63 were used to identify the crystallography of the nanoparticle using a copper filter. We evaluated the bonding of the nanocomposite and confirmed using Fourier transform infrared spectroscopy (FT-IR) with a Rayleigh-WQF-10 instrument in the range of 400–4000 cm^-1^.

## Results and Discussion

### FT-IR Analysis

IR-FT spectroscopy was used to identify functional groups and types of chemical bonds. As shown in Fig (1), the observed peak at 3448 cm ^-1^ is corresponded to OH water adsorption. Due to Nps high surface area and tend to adsorb water, the observed peak at 1596 cm ^-1^ is attributed to C-O bond. The observed peak at 618 cm ^-1^ and 435 cm ^-1^ corresponds to (M-O) Fe-O, metal-oxygen bond adsorptions.

**Figure (1):**
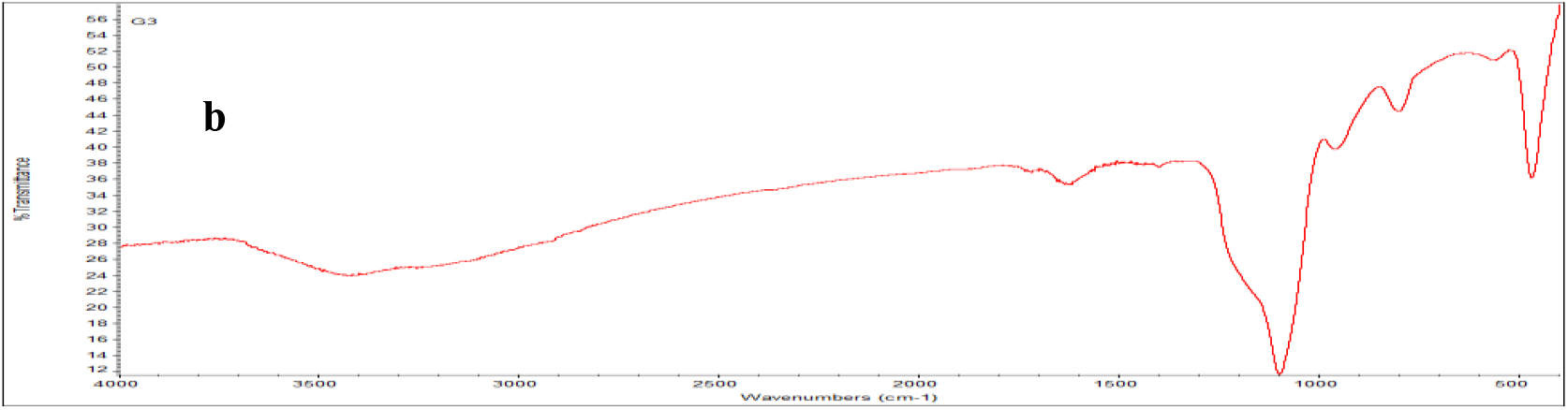
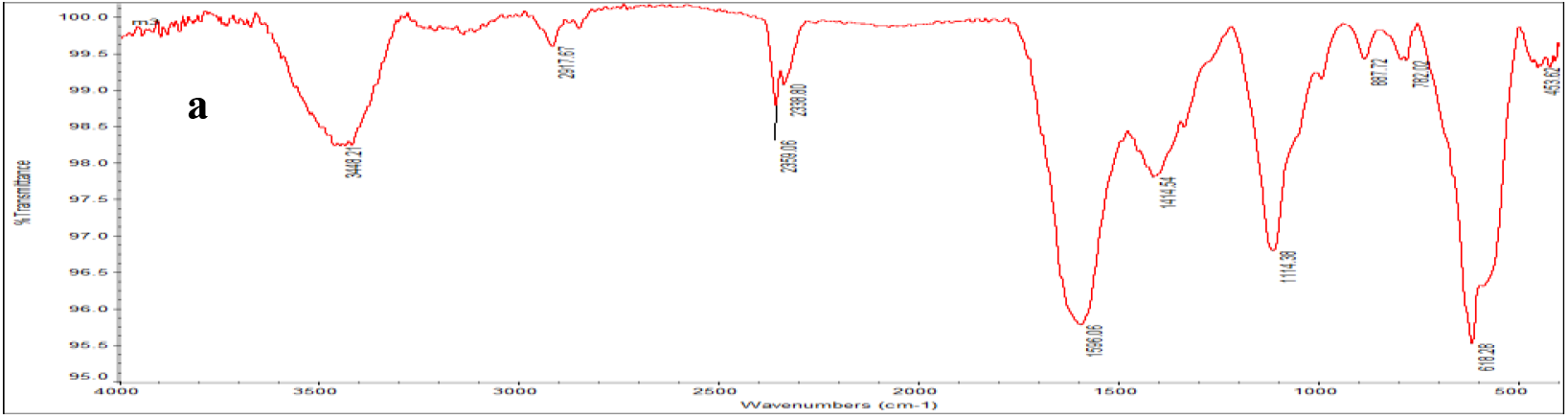
FT-IR Spectrum (**a** and **b** show FT-IR spectrum of synthesized iron oxide /graphene magnetic nanocomposite and graphene Nps, respectively)

### X-ray Diffraction Spectrum of Synthesized Magnetic Iron Oxide Graphene Nanocomposite

X-ray diffraction pattern of iron magnetic nano-oxide particles is shown in Fig (2). Peaks at angles 2θ = 32, 57, 46 and 63 were observed, indicating a crystalline nano-oxide structure. The found peaks confirm the rhombic structure of the material well, consistent with the research literature. XRD spectrum of nano-oxide Nps conforms to standard card (JCPDS 38-1479) and average particle size, determined using Scherrer^1^ formula, is approximately 60 nm, as shown in Fig(2). The average particle size can be calculated using the Debye-Scherrer equation provided in Eq (1)

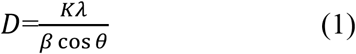

In equation above, *D* is the nanoparticle size, *K* is the aspect ratio or shape factor, λ is the wavelength, *β* is the Full width at half maximum (FWHM) and *θ* is the peak position on the horizontal axis of the diffraction pattern.

**Figure (2):**
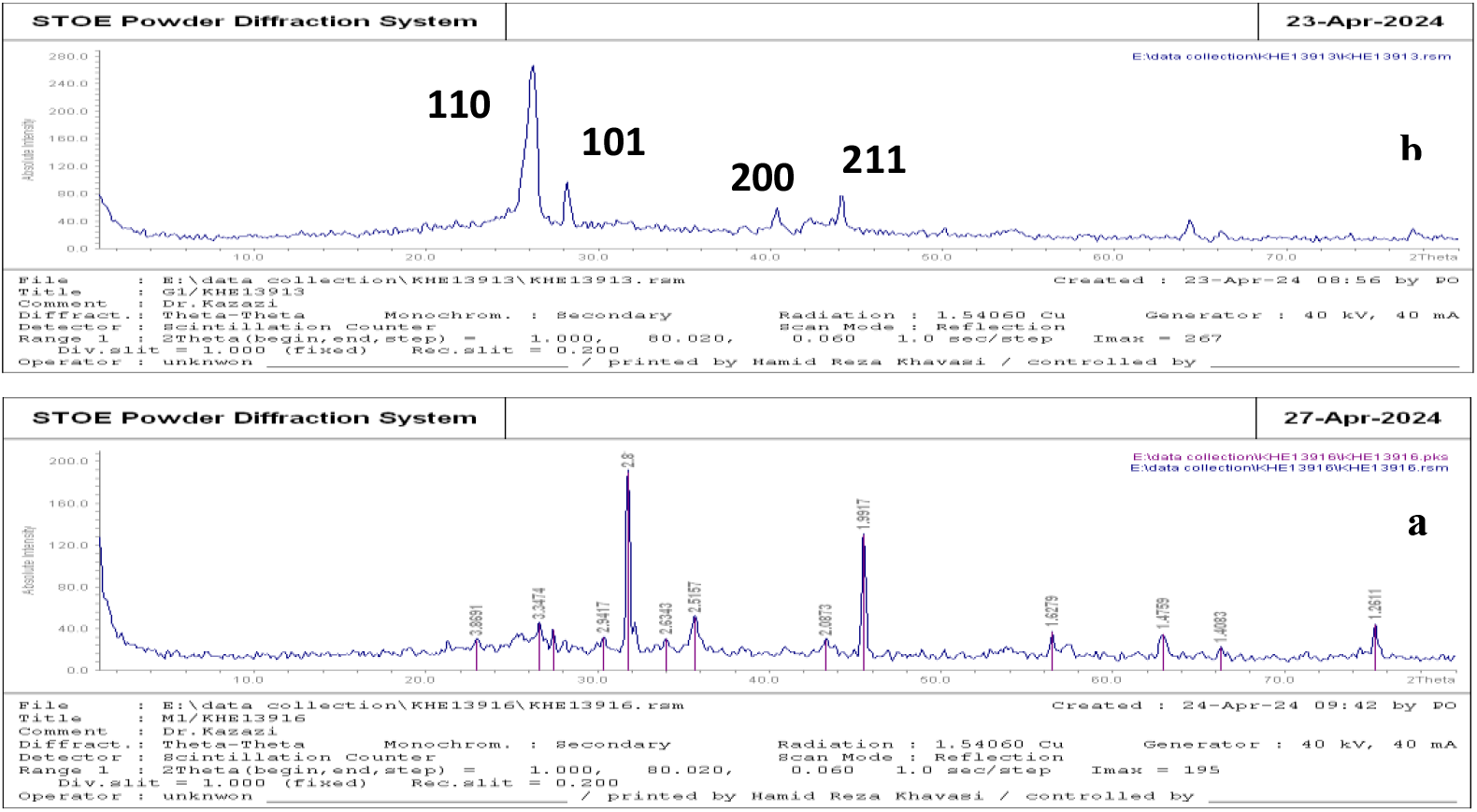
XRD spectrum (**a** and **b** show XRD spectrum of synthesized iron oxide /graphene magnetic nanocomposite and graphene Nps, respectively)

### Scanning Electron Microscopy (SEM) spectrum

Scanning Electron Microscopy (SEM) spectrum was utilized to investigate the morphology and distribution of Nps. SEM images were recorded at magnifications of 200, 500 nm, and 1 μ. The images illustrate Iron magnetic oxide Nps dispersed on Graphene sheets, with average particle sizes consistent with those obtained from XRD calculations. The results indicate that the iron oxide magnetic Nps are nano-sized and coated with graphene, as shown in Fig (3).

**Figure (3):**
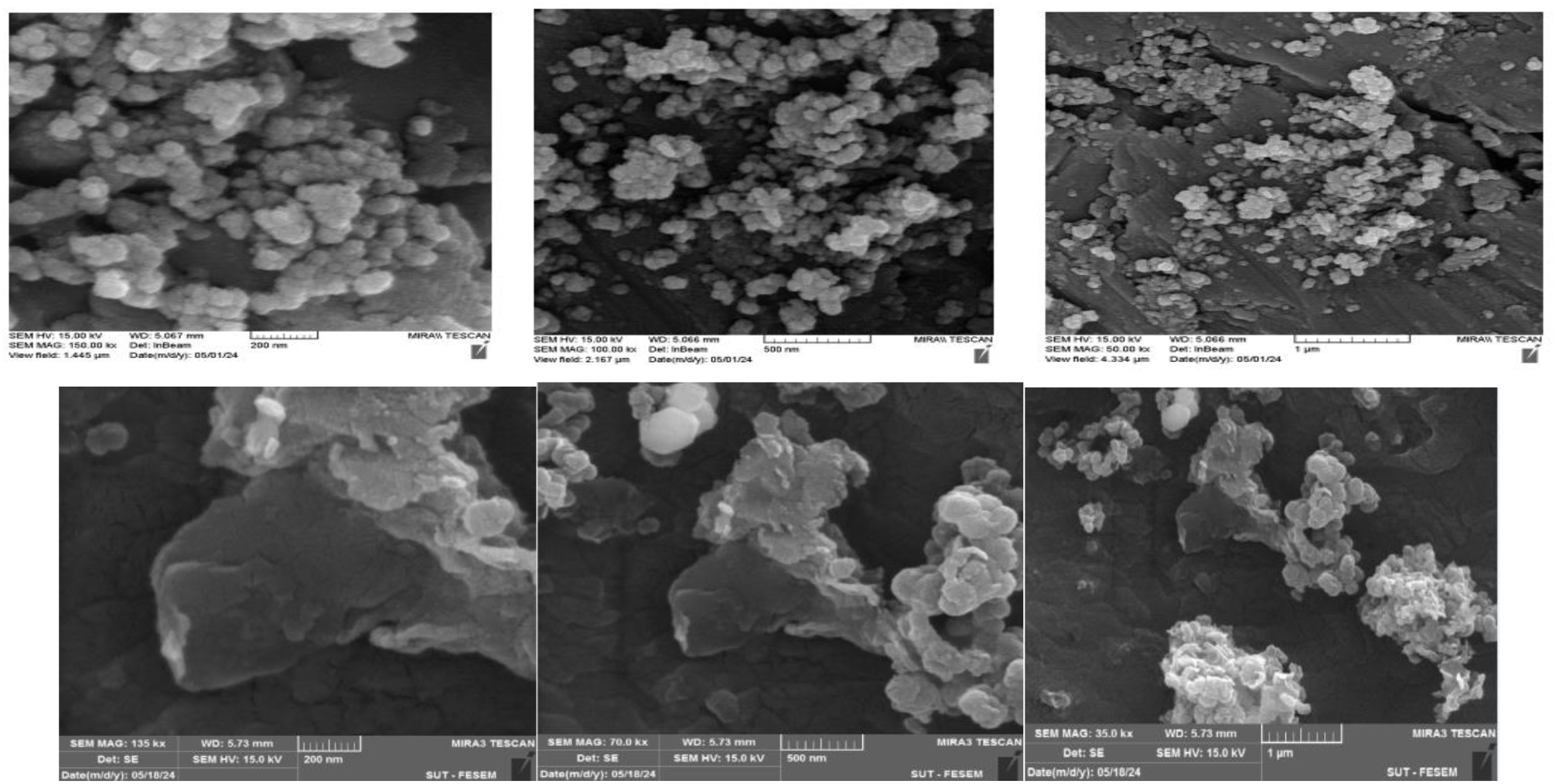
Scanning Electron Microscopy (SEM) spectrum (the top three images and the bottom three images show the SEM spectrum of iron oxide /graphene magnetic nanocomposite and Graphene Nps, respectively)

### Energy Dispersive X-ray Spectroscopy (EDX)

Energy Dispersive X-ray Spectroscopy (EDX) is an analytical method used for structural analysis or chemical characterization of a sample. This method relies on the interaction between an X-ray excitation source and a sample, determining the percentage of present elements in the sample, as shown in Fig (4).

**Figure (4):**
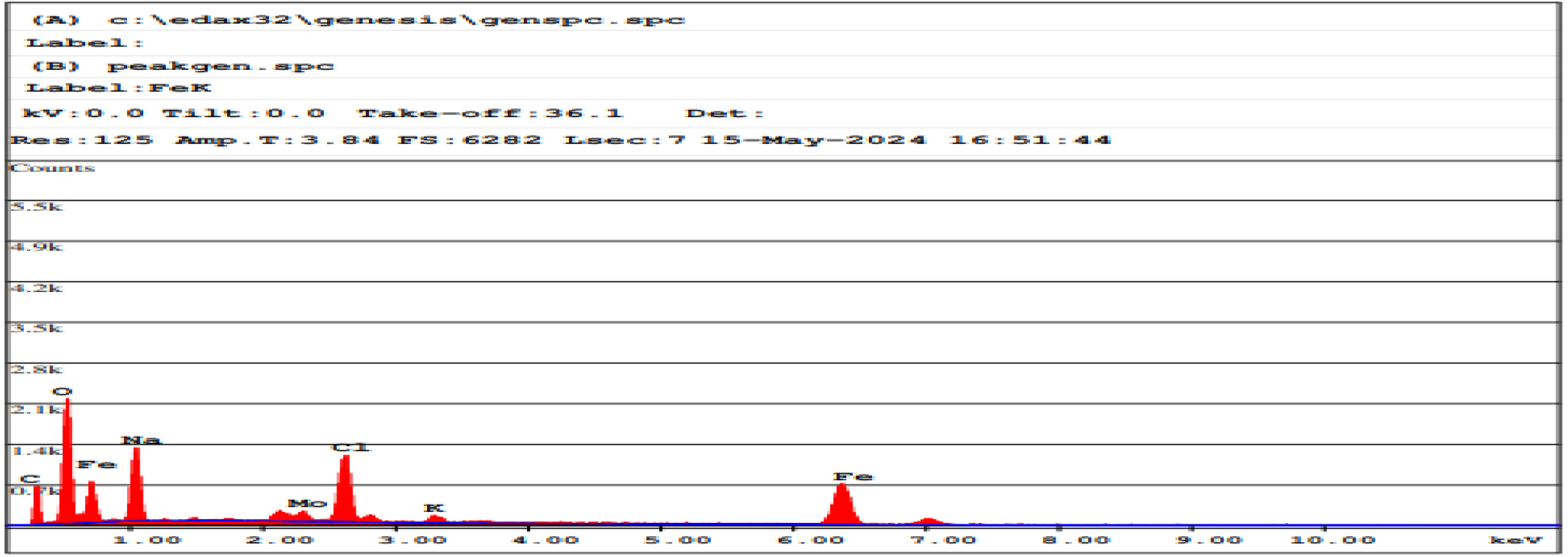
EDAX Spectrum of iron oxide magnetic Nps coated with graphene

According to the data, the tested sample contains 21% carbon, 36% oxygen, and 13% iron, table (1).

**Table (1):** Spectral data of magnetic iron oxide Nps coated with graphene (it should be noted that trace amounts of sodium, potassium and chlorine are false positives due to washing the material with distilled water)

| EDAX ZAF Quantification (Standardless) |  |  |  |  |  |  |
| --- | --- | --- | --- | --- | --- | --- |
| Element Normalized |  |  |  |  |  |  |
| SEC Table : Default |  |  |  |  |  |  |
| Element | Wt % | At % | K-Ratio | Z | A | F |
| C K | 9.95 | 21.26 | 0.0178 | 1.1010 | 0.1627 | 1.0004 |
| O K | 22.89 | 36.72 | 0.0769 | 1.0823 | 0.3098 | 1.0016 |
| FeL | 28.10 | 12.91 | 0.1205 | 0.9277 | 0.4619 | 1.0006 |
| NaK | 14.42 | 16.10 | 0.0349 | 1.0126 | 0.2390 | 1.0003 |
| MoL | 2.34 | 0.63 | 0.0178 | 0.8254 | 0.9161 | 1.0037 |
| ClK | 7.38 | 5.34 | 0.0604 | 0.9828 | 0.8293 | 1.0053 |
| K K | 0.89 | 0.58 | 0.0079 | 0.9870 | 0.8888 | 1.0130 |
| FeK | 14.04 | 6.45 | 0.1293 | 0.9198 | 1.0010 | 1.0000 |
| Total | 100.00 | 100.00 |  |  |  |  |

### Vibrating Sample Magnetometer (VSM) Magnetic Spectroscopy

VSM analysis is the primary method for studying the magnetic properties of materials. The result of this analysis is obtaining the hysteresis curve or remanence loop of materials, which can provide data such as coercivity, saturation magnetization and classification of material magnetism like ferromagnetic and superparamagnetic. Figure (5) shows the remanence loop of magnetic oxide Nps and Iron coated with graphite in fields ranging from -15000 to +15000 Oersted. The hysteresis curve demonstrates that the sample exhibits magnetic properties.

**Figure (5):**
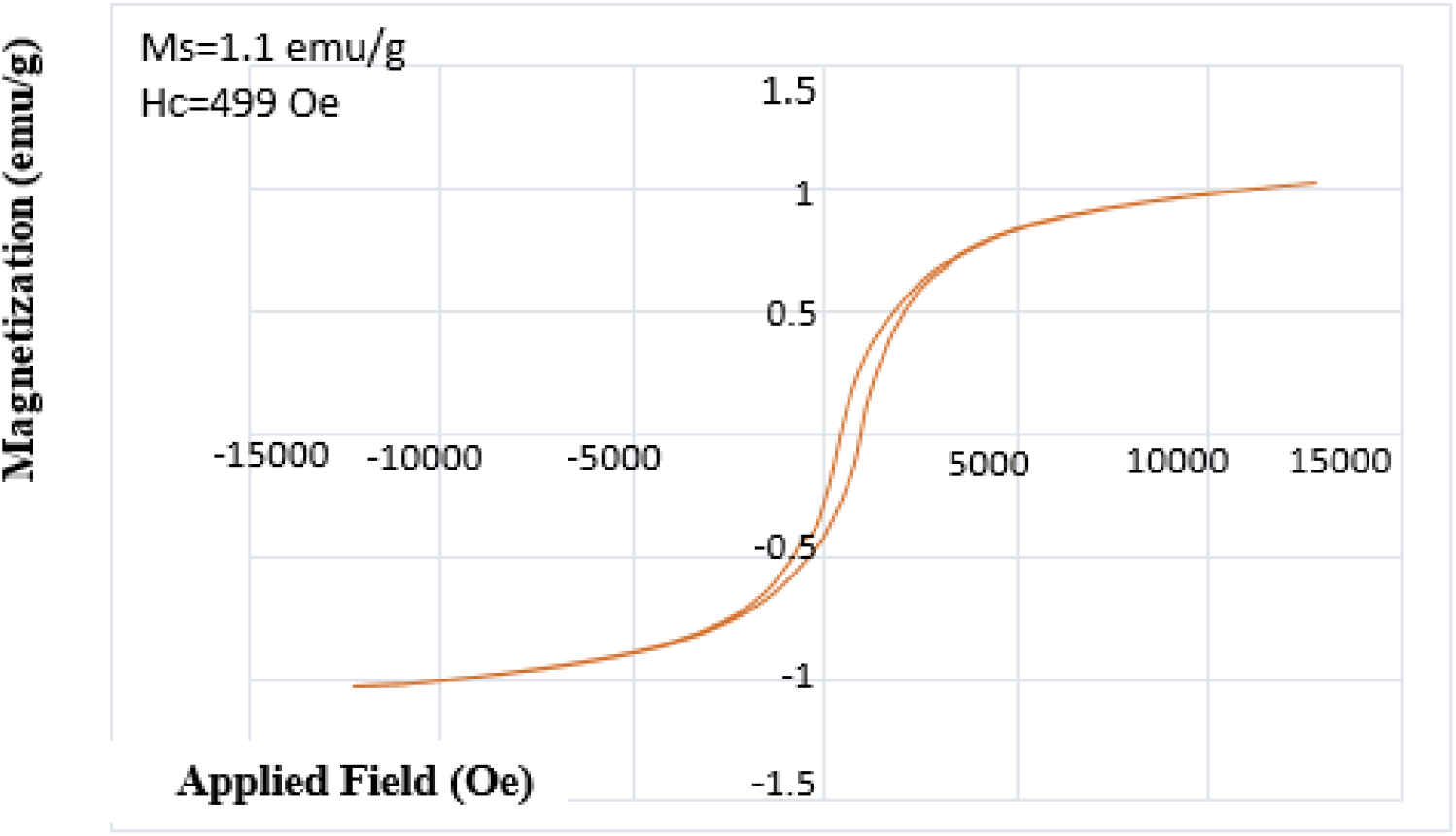
The Magnetization curve (VSM) of nanocomposite and its easy desorption from aqueous solutions by an external field

### Antibacterial Screening

Iron and Graphene ions inhibit the division of E. coli bacterial cells, damaging their cell walls and cellular contents, thereby preventing their growth and reproduction, as shown in Fig (6).

**Figure (6):**
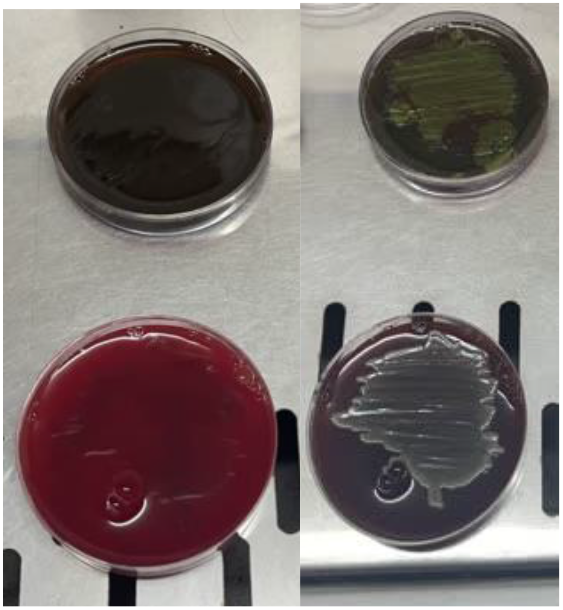
Culture of E.coli bacteria in Blood Agar medium

## Conclusion

In this study, we successfully synthesized iron oxide/graphene magnetic nanocomposite using green tea extract, known for its reducing and stabilizing agents. We investigated their antibacterial properties against E. coli bacteria. The results indicate a reduction in the proliferation of this bacterium. From this study, it is evident that the Fe_3_O_4_/G sample exhibits highly effective antibacterial properties, leading to inhibition of bacterial growth or reduction in growth specifically against E. coli. Additionally, the magnetic properties of this sample enable multiple uses.

## Footnotes

1 Debye Scherrer

